# Measuring excitation/inhibition balance through field potentials

**DOI:** 10.64898/2026.07.16.738654

**Authors:** Julia Rodriguez-Sanchez, Geoffrey W Diehl, Paul J Cunningham, Dimitris Pinotsis, Rick A Adams, A David Redish

## Abstract

Several electroencephalography-based metrics have been proposed to index excitation/inhibition balance (E/I). We used single-unit and local field potential recordings from rat medial prefrontal and orbitofrontal cortex and mouse visual cortex and hippocampus to evaluate four candidate metrics against empirically-measured E/I. While 1/f slope and gamma oscillations showed region and context dependencies, broadband (6-80 Hz) and high gamma (80-150 Hz) power consistently correlated with E/I, supporting their use as proxy measures. If at least 30 minutes’ data from medial prefrontal cortex were used per animal, broadband power could rank the animals by their E/I ratio, suggesting it could be used as an individual differences measure.

## Main

The balance between excitation and inhibition (E/I balance) is a fundamental property of cortical circuits. Disruptions in E/I balance have been implicated in neuropsychiatric disorders such as autism^1,2^ and schizophrenia^2,3^, and may underlie behavioural abnormalities and cognitive deficits^4,5^. Measuring E/I balance non-invasively remains challenging^6^, but several electroencephalography (EEG)-based metrics have been proposed as proxies, motivated by evidence that excitatory and inhibitory postsynaptic currents contribute to different components of EEG—as well as electrocorticogram (ECoG) and local field potential (LFP)—signals^7^.

One purported proxy of E/I balance is gamma-band oscillations (30-80 Hz), which typically emerge from coordinated interactions between principal cells and fast-spiking GABAergic interneurons^8^. Inhibition and stimulation of parvalbumin (PV)-expressing interneurons has been shown to suppress and increase gamma oscillations, respectively^9,10^, suggesting reduced gamma-band power may reflect reduced PV inhibition (although other interneuron subtypes have also been implicated^11,12^).

More recently, computational modelling work by Gao et al.^13^ suggested that the aperiodic, or “1/f”, component of the power spectrum reflects the balance between (faster) AMPA and (slower) GABA_A_ synaptic conductances, with steeper slopes reflecting greater inhibition. This was validated empirically in rat hippocampal CA1^13^. In humans and macaques, steeper 1/f slopes reflected drug-induced decreases in the E/I ratio^13,14^.

Although widely used as proxies of E/I balance, these measures have rarely been evaluated together (cf.^15^), and their relationship to the activity of excitatory and inhibitory cells has not been assessed systematically. A potential problem with pharmacological validation is that GABAergic drugs can reduce PV firing^16–18^ and so could paradoxically increase E/I activity ratio whilst reducing E/I conductance ratio. This makes it crucial to assess E/I balance measures under physiological conditions.

To address this, we leveraged large-scale silicon probe recordings from medial prefrontal cortex (mPFC) and orbitofrontal cortex (OFC) of rats performing the naturalistic decision-making Restaurant Row task^19^, and Neuropixels recordings from mouse primary visual cortex (V1) and hippocampal CA1 during spontaneous and visual-evoked activity^20^. We extracted excitatory and inhibitory firing rates alongside LFP metrics, and investigated the relationship between purported electrophysiological proxies of E/I balance and empirically measured E/I ratios across species, cortical regions and behavioural states. We also assessed broadband (6-80 Hz) and high-gamma (80-150 Hz) power, which have been shown to correlate with single-neuron firing rates in humans^21,22^ and macaque visual cortex^23^, respectively, and with E/I ratio in rats^24^.

A total of 11057 single units were recorded across species and regions, including 2898 from rat mPFC, 5595 from rat OFC, 862 from mouse V1, and 1702 from mouse CA1. Units were classified as excitatory or inhibitory using two approaches: spike width, with units below 0.4 ms classified as putative interneurons and those above as putative principal cells (**Fig. 1a and 1c**), and spike cross-correlation histograms (CCH) for each pair of simultaneously recorded cells, with presynaptic cells classified based on connection sign^25^ (**Fig. 1b and 1d**). Spike width-based and CCH-based cell classifications were largely consistent in mPFC and V1 (>80% correspondence) but diverged in CA1, where synchronous population activity makes it difficult to identify putative monosynaptic connections, and OFC, where width-based classification was unreliable due to low spike waveform quality (**Fig. 1e**). We used spike width-based classification for mPFC, V1 and CA1, and CCH-based classification for OFC, with analyses using the alternative measure shown in **Supplementary Fig. 2**.

**Figure 1.**
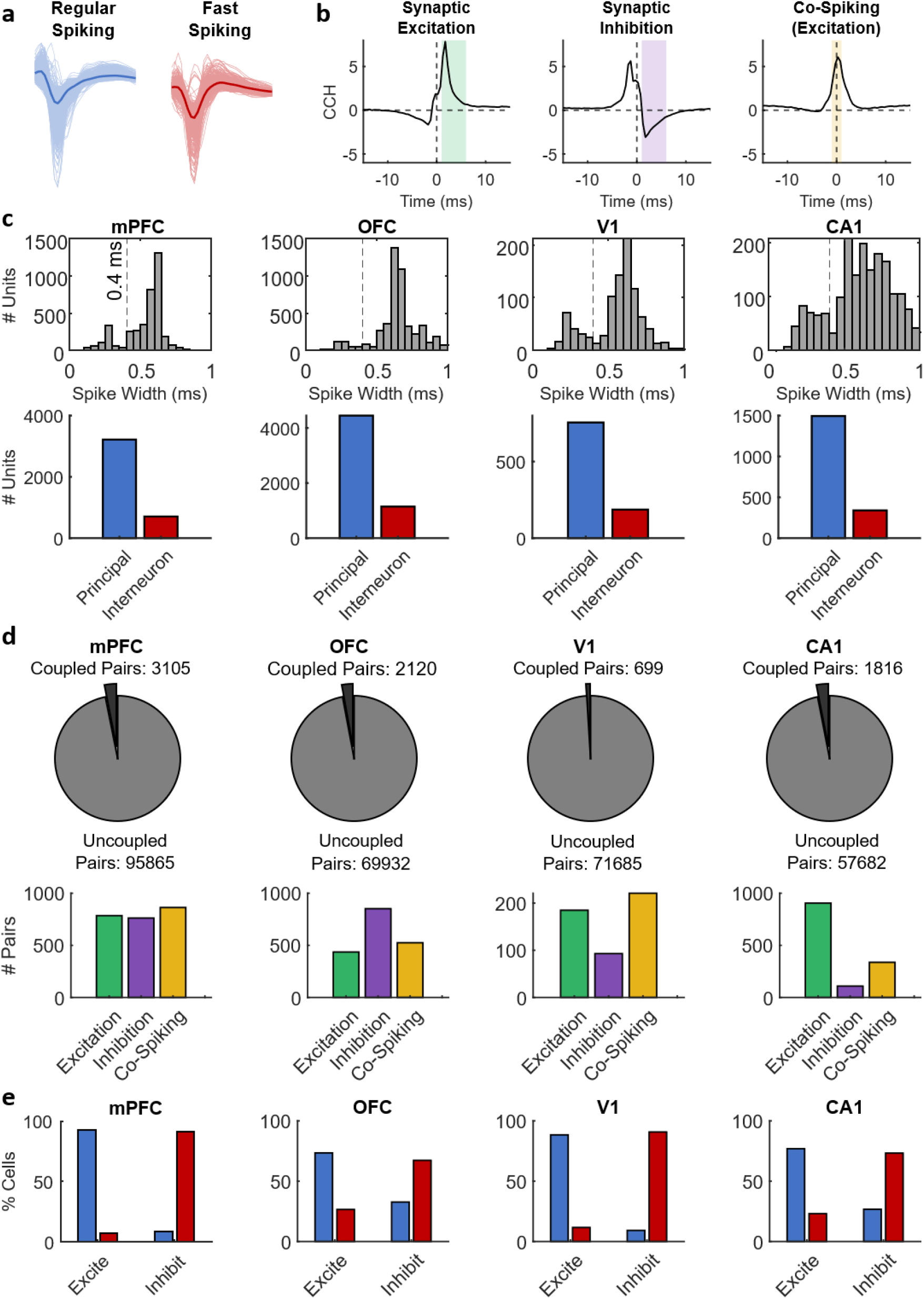
Cell type classification. **(a)** Putative regular-spiking principal cells and fast-spiking interneurons were identified based on the spike-width distributions, illustrated here with average waveforms recorded from medial prefrontal cortex (mPFC). **(b)** Putative excitatory and inhibitory synaptic connections were identified using high-resolution spike cross-correlation histograms (CCH) and classified as synaptic excitation (positive deflection between 1-6 ms), synaptic inhibition (negative deflection between 1-6 ms) or co-spiking (deflection at <1 ms), illustrated here with z-scored CCH responses for those cell pairs with identified synaptic connections in mPFC. Only synaptic excitation and synaptic inhibition were analysed. **(c)** Top: Spike width distributions for units recorded in mPFC, orbitofrontal cortex (OFC), primary visual cortex (V1) and hippocampal CA1. The dashed line at 0.4 ms separates putative interneurons (narrow spikes) from putative principal cells (broad spikes). Bottom: Number of principal cells (blue) and interneurons (red) identified in each region. **(d)** Top: Proportion of coupled and uncoupled cell pairs identified via CCH analysis across mPFC, OFC, V1 and CA1. Bottom: Breakdown of coupled pairs by connection type in each region. **(e)** Proportion of excitatory or inhibitory synaptically-coupled projection cells that were identified as principal cells (blue) or interneurons (red).

From these putative cell classifications, E/I ratios were computed as the ratio of mean excitatory to inhibitory firing rates across 512 ms windows (**Supplementary Fig. 1a**)and correlated against the four LFP metrics quantified for the same time bins: 1/f slope (30-50 Hz), oscillatory gamma power (30-80 Hz), overall broadband power (6-80 Hz), and high gamma power (80-150 Hz) (**Supplementary Fig. 1b**). Additional analyses investigated total (i.e., not aperiodic-corrected) power in the 30-80 Hz range (**Supplementary Fig. 3**).

We first assessed the relationship between E/I ratio and LFP metrics during spontaneous activity (**Fig. 2**). Consistent with computational predictions^13^, 1/f slope correlated negatively with E/I ratio in hippocampal CA1 (Mann-Whitney U test; z=-2.47, p=.014, d’=0.44; **Fig. 2a**). However, no significant relationship between 1/f slope and E/I ratio was observed in OFC (z=1.16, p=.244, d’=0.12) or V1 (z=-0.81, p=.420, d’=0.09), and there was a weak *positive* correlation in mPFC (z=2.77, p=.006, d’=0.21; **Fig. 2a**). Oscillatory gamma power correlated negatively with E/I ratio in mPFC (z=-2.59, p=.010, d’=0.23) and CA1 (z=-4.45, p<.001, d’=0.79), but not in OFC (z=1.69, p=.092, d’=0.19) or V1 (z=1.50, p=.133, d’=0.25; **Fig. 2c**); in contrast, total 30-80 Hz power correlated negatively with E/I ratio across all four regions (**Supplementary Fig. 3a-b**). Broadband (**Fig. 2b**) and high gamma power (**Fig. 2d**) correlated negatively with E/I ratio across mPFC (Broadband: z=-2.73, p=.006, d’=0.29; High Gamma: z=-4.62, p<.001, d’=0.52), OFC (Broadband: z=-4.02, p<.001, d’=0.44; High gamma: z=-6.99, p<.001, d’=0.74), V1 (Broadband: z=-3.99, p<.001, d’=0.70; High Gamma: z=-6.87, p<.001, d’=1.42) and CA1 (Broadband: z=-4.94, p<.001, d’=0.71; High Gamma: z=-8.35, p<.001, d’=1.85). These negative correlations arose despite broadband and high gamma power correlating positively with both excitatory and inhibitory firing rates (**Supplementary Fig. 4**). Firing rate variability (coefficient of variation) was higher for putative inhibitory than putative excitatory cells in mPFC, V1 and CA1 (p<.05) but not in OFC (**Fig. 2e**).

**Figure 2.**
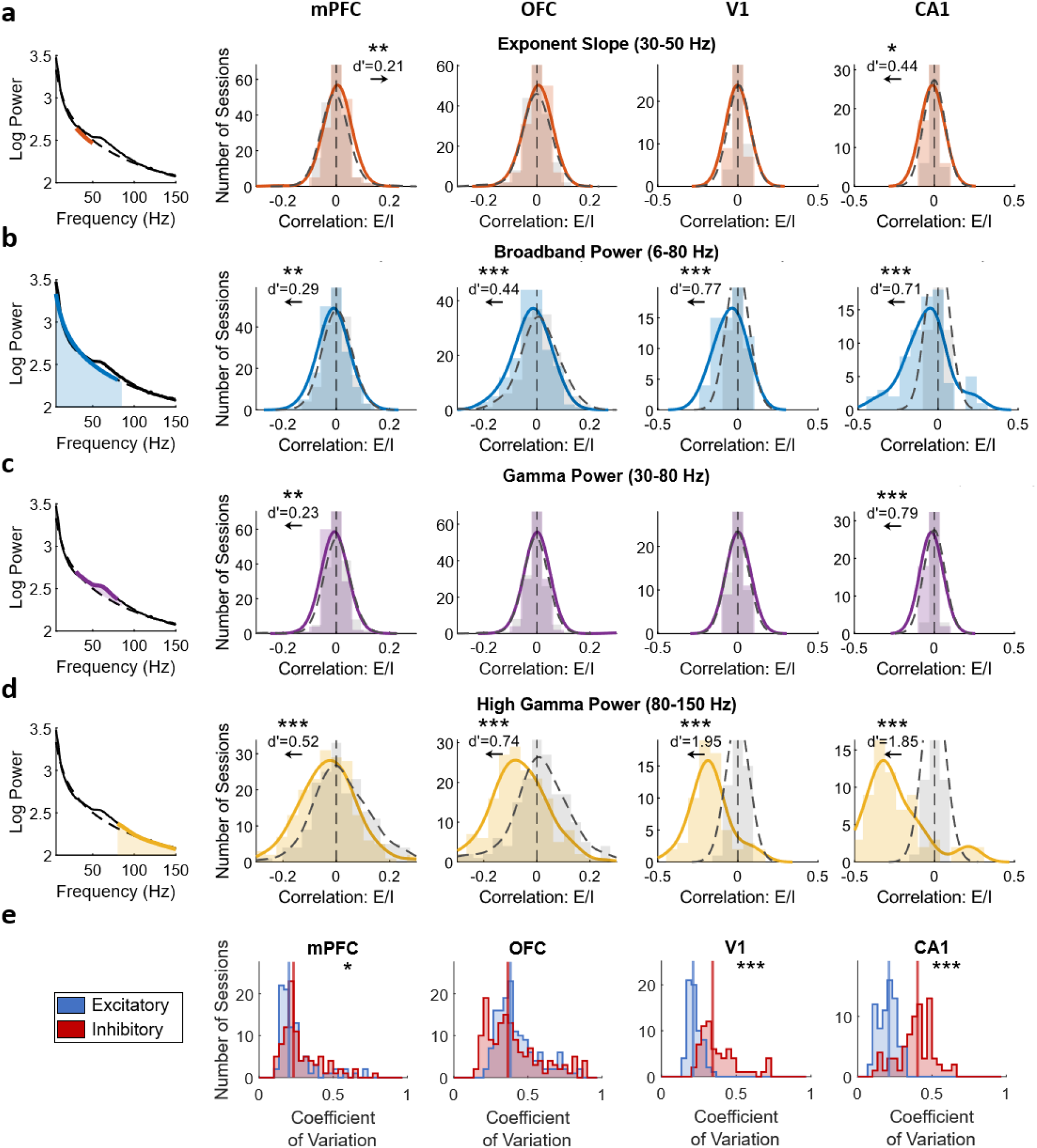
Local field potential metrics track excitation/inhibition balance across regions during spontaneous (non-task-related) activity. **(a-d)** Schematic illustration of each local field potential (LFP) metric (left) and the corresponding distributions of correlation coefficients between that metric and the E/I ratio across medial prefrontal cortex (mPFC), orbitofrontal cortex (OFC), primary visual cortex (V1), and hippocampal CA1 (right, ordered left to right). Coloured histograms (solid curves) show real data; grey dashed curves show a shuffled null distribution. Note that the x-axis scale for mPFC and OFC differs from V1 and CA1. **p<.01, ***p<.001. **(e)** Distribution of coefficient of variation of excitatory (blue) and inhibitory (red) firing rates across mPFC, OFC, V1 and CA1. Coefficient of variation was computed across recording days for mPFC and OFC, and across temporal chunks within a single recording session for V1 and CA1. *p<.05, ***p<.001.

To further investigate whether E/I-LFP relationships were behaviourally context-dependent, we next analysed rat mPFC and OFC recordings during the Restaurant Row task (**Fig. 3a**) and mouse V1 recordings during presentation of drifting visual gratings (**Fig. 3b**). In mPFC, the positive correlation between E/I ratio and 1/f slope (Mann-Whitney U test; z=1.98, p=.048, d’=0.29) and negative correlations with broadband (z=-4.54, p<.001, d’=0.54) and high gamma (z=-5.37, p<.001, d’=0.72) power were also observed during task engagement, but the negative correlation between E/I ratio and oscillatory gamma power observed during spontaneous activity was absent during the decision-making task (p>.05; **Fig. 3c**). In OFC, negative correlations between E/I ratio and broadband (z=-3.33, p<.001, d’=0.66) and high gamma (z=-4.74, p<.001, d’=0.67) power were also observed during task engagement, and there was a positive correlation with 1/f slope (z=3.15, p=.002, d’=0.51) that was absent during spontaneous activity. In V1, E/I ratio correlated negatively with broadband power (z=-3.27, p=.001, d’=0.52) and high gamma power (z=-2.86, p=.004, d’=0.53) during presentation of a visual grating (**Fig. 3d**), consistent with the spontaneous results.

**Figure 3.**
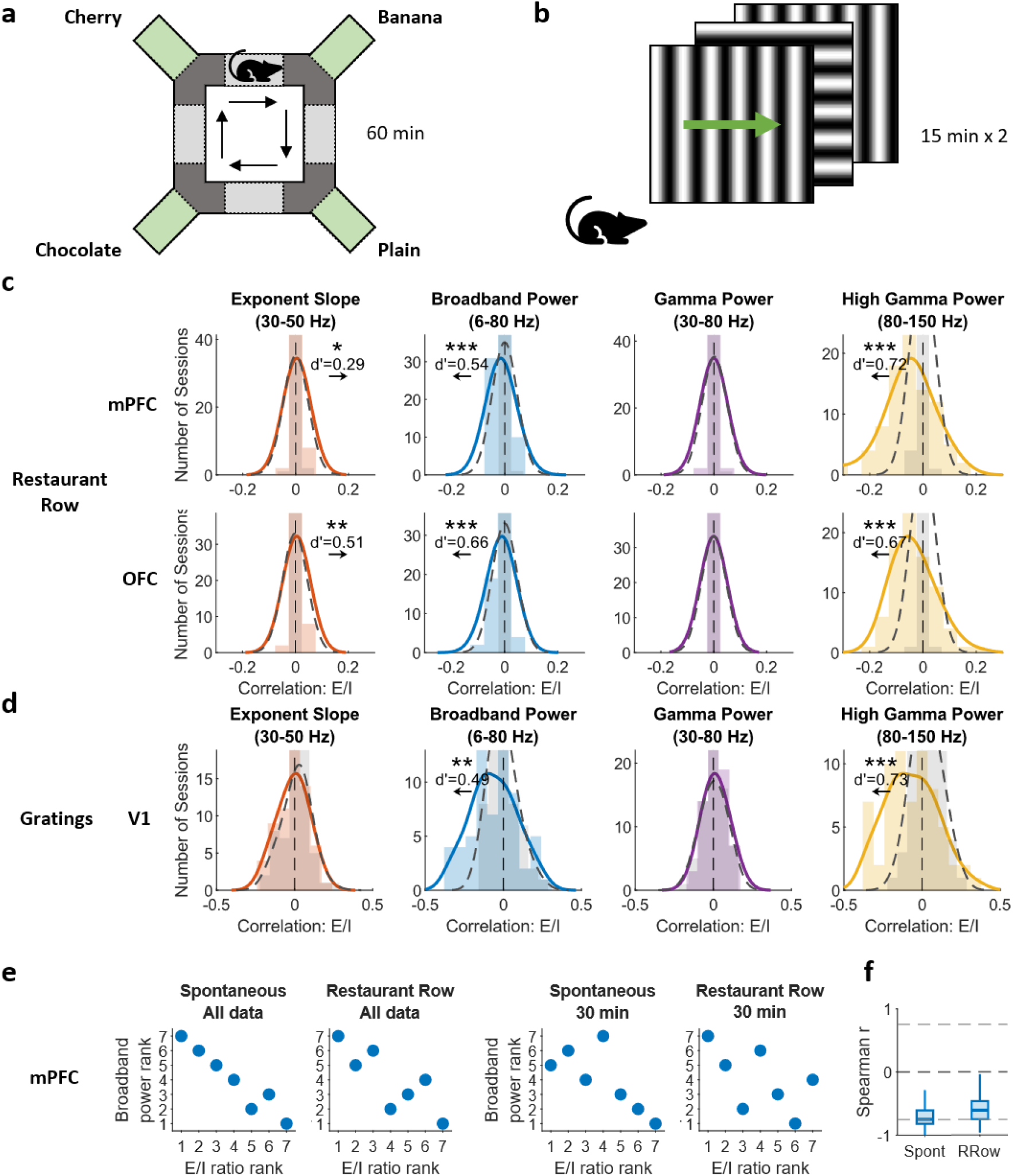
Local field potential metrics track excitation/inhibition balance and can rank animals based on excitation/inhibition ratio during task- and stimulus-driven activity. **(a)** Schematic of the Restaurant Row (RRow) decision-making task. Rats ran around a circular maze with four reward sites (“restaurants”) offering different food flavours over a 60-minute session. **(b)** Schematic of the visual stimulation task. Mice were presented with drifting gratings over two sessions of 15 minutes each. **(c-d)** Distributions of correlation coefficients between local field potential (LFP) spectral features (aperiodic exponent, broadband power, gamma power, high-gamma power) and the excitation/inhibition ratio in medial prefrontal cortex (mPFC), orbitofrontal cortex (OFC) and primary visual cortex (V1). Coloured histograms (solid curves) show real data; grey dashed curves show a shuffled null distribution. *p<.05, **p<.01, ***p<.001. **(e)** Rank-order correspondence between animals’ mean E/I ratio and broadband power (6-80 Hz) in mPFC. Each dot represents one animal. Animals are ranked from lowest to highest E/I ratio on the x-axis, and ranked from lowest to highest broadband power on the y-axis. Left panels show rankings using all available data; right panels show rankings using 30 minutes of data per animal (10 minutes per day, averaged across 3 days). **(f)** Distribution of Spearman correlation coefficients between mPFC E/I ratio and broadband power rank across 1,000 subsampling iterations. Dashed gray lines indicate the threshold for statistical significance at p<.05 (two-tailed).

To assess whether broadband and high gamma power also track between-animal differences in E/I balance, we ranked animals by their mean E/I ratio and tested whether this correlated with LFP metric rankings. In mPFC, broadband power reflected the animal-level E/I ranking both during spontaneous activity (r=-0.96, p<.001) and during the Restaurant Row task (r=-0.82, p=.023; **Fig. 3e**). High gamma power showed a similar non-significant trend (**Supplementary Fig. 5**). No significant relationship was observed in OFC, V1 or CA1 (**Supplementary Fig. 5**). To test if this was driven by differences in recording duration, we repeated the mPFC broadband power analysis using only 30 minutes of data (1,000 iterations; spontaneous: median r=-0.71, Restaurant Row: median r=-0.59; **Fig 3e-f**).

This study systematically evaluated purported electrophysiological proxies of E/I balance. Oscillatory gamma power (30-80 Hz) and 1/f slope (30-50 Hz)—often used to estimate E/I balance non-invasively^8,13,26–28^—showed markedly inconsistent relationships with empirically measured E/I ratio across brain regions, challenging their generalisability as E/I proxies. In contrast, broadband (6-80 Hz) and high gamma (80-150 Hz) power consistently correlated negatively with E/I ratio across species, regions and behavioural contexts.

The 1/f slope correlated negatively with E/I ratio in hippocampal CA1, positively in mPFC, and not in OFC or V1. This regional heterogeneity suggests that prior validations using LFP recordings in hippocampal CA1^13^ do not generalise to other structures. This may reflect various issues: the Gao et al.^13^ model included only AMPA and GABA receptors, omitting NMDA receptors which have substantially longer time constants, and it defines E/I as a ratio of conductances, not firing rates. Moreover, modelling work shows that the direction of the E/I-slope relationship can reverse depending on leak conductance^29^. At the circuit level, the direction of the slope change depends on which interneuron subtype is engaged: optogenetic suppression of somatostatin interneurons in mouse V1 produced the expected flattening, but suppressing PV interneurons increased principal cell spiking yet steepened the 1/f slope^30^. In line with this parameter-dependence, lorazepam both flattened and steepened 1/f slope in different cortical areas in humans, with the greatest flattening in regions with highest GABA_A_ receptor density^31^.

The relationship between oscillatory gamma power and E/I ratio was also region- and context-dependent: oscillatory gamma power correlated negatively with E/I in mPFC and CA1, but not in OFC or V1, and the negative correlation in mPFC was absent during engagement in a decision-making task. However, total (i.e., not aperiodic-corrected) spectral power in the 30-80 Hz range did show consistent correlations with E/I ratio. The correlation between spiking and LFP power has been shown to flip from negative between 1-30 Hz to positive between 30-100 Hz^22^, so broadband power increases due to changes in population firing rate may manifest as increases in the gamma band. Apparent associations between gamma power and E/I ratio may therefore reflect aperiodic broadband activity rather than narrowband oscillations if oscillatory and aperiodic spectral components are not separated before analysis^32^.

Broadband and high gamma power reflected excitatory and inhibitory firing rates across all four regions, consistent with previous findings in humans^21^ suggesting that LFP signals reflect the sum of postsynaptic currents, such that increased population firing shifts the broadband spectrum upwards^30^. In contrast, narrowband oscillatory activity exhibits more variable relationships with firing, depending on local network synchrony and neuron-specific dynamics^33^—and indeed we saw a positive relationship between oscillatory gamma and interneuron firing in mPFC and CA1 but not OFC or V1. Under this framework, broadband and high gamma power likely track overall population firing activity, and the negative correlation with E/I ratio likely arises because of greater variability in inhibitory relative to excitatory firing. Importantly, this relationship held not only across sessions but also across animals: in mPFC, individual differences in broadband and high gamma power reflected individual differences in E/I ratio.

Notably, we identified putative excitatory and inhibitory cells based on either waveform characteristics or CCH-based synaptic connectivity. While a widely used approach, waveform-based classification entails inherent misclassification errors^34^ and is biased towards detecting fast-spiking PV interneurons—although, of all interneurons, PV cells probably contribute most to inhibition (given they synapse near the soma with high conductance and high connection probability). CCH-based classification is limited to cells forming detectable synaptic connections with co-recorded units, which may not be representative of the broader population. Nevertheless, both approaches produced broadly consistent negative correlations between broadband and high gamma power and E/I.

Together, these results indicate that broadband and high gamma power may track E/I balance across brain regions and behavioural contexts, whereas 1/f slope and oscillatory gamma are unlikely to provide robust correlates. Given the growing interest in EEG-based proxies of E/I balance^13,28,35^, these findings highlight the importance of empirically validating E/I metrics before translating them to human studies, and suggest that broadband power, which can be measured non-invasively using EEG, and high gamma power, which can be measured with ECoG and intracranial LFP, may be more promising tools for tracking E/I balance in clinical settings.

## Online Methods

### Animals

Data were drawn from three sources: a previously published dataset comprising recordings from a cohort of eight adult Fisher-Brown Norway F1 hybrid rats (4M, 4F; aged 9-15 months; cohort A)^19^, a previously published dataset comprising recordings from a cohort of seven adult Fisher-Brown Norway rats (4F, 3M; aged 8-18 months; cohort B)^36^, and a publicly available dataset of 14 adult wild-type C57BL/6J mice (11M, 3F; aged P108-P144) from the Allen Brain Observatory Visual Coding Neuropixels dataset^20^. One rat (R506) from cohort A was excluded due to differences in recording methodology. One rat (R676) from cohort B was excluded due to low cell yields. Two mice (771990200, 847657808) were excluded due to missing data segments. This resulted in effective sample sizes of n=7 rats (3M, 4F) in cohort A, n=6 (3M, 3F) rats in cohort B, and n=12 mice (10M, 2F). All analyses presented here are novel and extend beyond those reported in the original publications.

Rat experiments were approved by the University of Minnesota Institutional Animal Care and Use Committee (IACUC) and were performed in accordance with NIH guidelines. The Allen dataset was accessed under its open data license and in accordance with the Allen Institute’s ethical standards.

### Task

Rats performed the decision-making Restaurant Row task over a 70-minute session, as detailed previously^19,36^. Each rat completed one session per day over 14 days. One session (from R535) was excluded because neural data could not be synchronised to behaviour. Sessions comprised a 60-minute behavioural task, during which rats foraged across four reward sites (“restaurants”) offering different food flavours, and two 5-minute periods of quiet rest away from the task before and after. During the task, the frequency of an auditory cue signalled the delay (1-30 s) to obtain the reward at each site, and rats decided whether to accept the offer and wait or skip to the next site. If the offer was accepted, the delay counted down via tone cues of decreasing frequency until reward delivery or the rat quitting. Task and rest periods were analysed separately.

Mice were head-fixed and passively viewed sequences of drifting gratings, natural movies, and dot motion stimuli over one ∼3-hour session, as detailed previously^20^. Analyses in the present study were restricted to two 15-minute blocks of drifting gratings and a 30-minute block of spontaneous activity. Drifting gratings were shown with a spatial frequency of 0.04 cycles per degree, two contrasts (0.1, 0.8), four orientations (0°, 45°, 90°, 135°) and a temporal frequency of 2 Hz, with 75 repeats. Spontaneous activity was recorded during presentation of a mean-luminance blank screen. Drifting grating and spontaneous activity periods were analysed separately. Each block was segmented into non-overlapping 5-minute analysis windows (“pseudo-sessions”) to provide multiple within-animal observations and enable comparable analyses across species.

### Recordings

All datasets included simultaneous single-unit spiking activity and LFPs. Rat recordings were obtained from mPFC and OFC using 256 channel Intan RHD recording systems connected to 64-channel Intan RHD headstages. Signals were sampled at 30 kHz. Single units were identified using Kilosort v2.0 (https://github.com/MouseLand/Kilosort)^37,38^ following median reference subtraction and high-pass filtering at 600 Hz. LFPs were extracted by downsampling to 2000 Hz and low-pass filtering at 800 Hz.

Mouse recordings were obtained from V1 and CA1 using six Neuropixels probes. The Allen Institute’s Neuropixels pipeline has been described in detail in Siegle et al.^20^. Signals were sampled at 30 kHz (spike band, high-pass filtered at 500 Hz) and 2500 Hz (LFP band, low-pass filtered at 1000 Hz). Single units were identified using Kilosort v2.0^37,38^ following median reference subtraction (within and across channels), high-pass filtering at 150 Hz and whitening in blocks of 32 channels. LFPs were extracted by downsampling to 1250 Hz and removing three out of every four channels.

### Software

All analyses were run in MATLAB (version R2018b; The MathWorks Inc., Natick, MA, USA; https://www.mathworks.com). Data from the Allen Brain Observatory Visual Coding Neuropixels project were accessed and formatted using a Jupyter Notebook for compatibility with the analysis pipeline.

### Cell class designations

Single units were classified as putative “principal cells” or “interneurons” based on spike width (i.e., the difference between the cell’s average waveform peak and trough), with units below 0.4 ms classified as interneurons and those above classified as principal cells (**Fig. 1a and 1c**). For the Restaurant Row dataset, mean spike waveforms were taken from the channel with the largest amplitude, assumed to be closest to the cell body. As a quality control criterion, units with spike widths exceeding 1 ms were excluded prior to classification (692 cells in OFC, 0 cells in mPFC), as widths of this magnitude likely reflect incorrect channel selection. For the Allen Brain Observatory dataset, spike widths were taken from the dataset metadata^20^. This yielded 2301 principal cells and 597 interneurons in mPFC, 4449 principal cells and 1146 interneurons in OFC, 701 principal cells and 161 interneurons in V1, and 1407 principal cells and 295 interneurons in CA1 (**Fig. 1c**).

### Synaptic coupling

Putative excitatory and inhibitory synaptic connections were identified using high-resolution spike cross-correlation histograms (CCHs) computed for each pair of simultaneously recorded cells, as described previously^25^. Identified coupling relationships were classified as synaptic excitation (positive deflection between 1-6 ms), synaptic inhibition (negative deflection between 1-6 ms) or co-spiking (deflection at <1 ms; **Fig. 1b**).

### Measures of excitation/inhibition

Firing rates were computed for all units in 512 ms bins with no overlap. Excitation and inhibition were defined as either (1) the mean firing rate of all identified principal cells and interneurons, respectively, or (2) the mean firing rate of all presynaptic cells with an identified synaptic excitatory or inhibitory connection, respectively. The E/I ratio was calculated for each bin as excitation divided by inhibition (**Supplementary Fig. 1**). Only recording sessions containing at least one putative excitatory and one putative inhibitory cell were included.

### Local field potential metrics

Up to three LFP channels per probe were randomly selected for analysis, excluding any channels where more than 20% of samples were saturated. Spectral power was estimated using short-time Fourier transform (*spectrogram* function in MATLAB), computed in 512 ms windows with no overlap (**Supplementary Fig. 1d**). For the gratings stimulus analysis, LFP spectra and firing rates were computed within individual 512-ms windows locked to stimulus onset, rather than across consecutive sliding windows, in order to restrict the analysis to periods of active visual stimulation.

A 1024-sample Hamming window was used for rat data (2000 Hz sampling rate) and a 640-sample window for mouse data (1250 Hz). Power was computed from 6-150 Hz in 0.5-Hz increments, and each power spectrum was parameterised using the MATLAB implementation of the FOOOF algorithm, which decomposes the power spectrum into an aperiodic component and superimposed periodic (oscillatory) peaks. The aperiodic component was modelled in two ways: using a “fixed” model fitted between 30-50 Hz to extract the 1/f exponent, and using a “knee” model fitted between 6-80 Hz to extract gamma and broadband power. If the knee parameter was negative, the knee fit was replaced by a fixed fit.

The 1/f slope was quantified as the FOOOF-derived aperiodic exponent between 30-50 Hz. Broadband power was quantified as the mean power of the fitted aperiodic component between 6-50 Hz. Gamma power was quantified as the mean power of FOOOF-identified oscillatory peaks between 30-80 Hz (note that we focused on rhythmic activity, which we explicitly separated from the aperiodic component, but see **Supplementary Fig. 3** for additional analyses using mean raw log10 power between 30-80 Hz). High gamma power was defined as the mean raw log10 power between 80-150 Hz.

### Statistics

For each session, Pearson correlations between each LFP metric and the E/I ratio time series were computed, then aggregated across LFP channels and probes to yield the median correlation per session. Correlations between LFP metrics and E/I ratio were assessed using Mann-Whitney U tests against a null distribution generated by shuffling cell inter-spike intervals (ISIs). For each cell, ISIs were randomly permuted once to preserve the overall firing rate while disrupting relationships with LFP and other cells’ spiking. Shuffled spike trains were used to recompute E/I ratios, which were then correlated with LFP metrics and again aggregated by median, yielding one null correlation value per session. The distribution of observed correlations across sessions was then compared against the distribution of null correlations to assess statistical significance. We report effect sizes (d’) for two sample comparisons.

All analyses were repeated using a single LFP channel per probe to reduce the potential influence of correlated LFP channels (**Supplementary Fig. 6**). To dissociate the contributions of excitatory and inhibitory activity, the same analysis was repeated correlating each LFP metric with mean excitatory and mean inhibitory firing rates separately, instead of the E/I ratio. To assess generalisation across animals, animals were additionally ranked by their mean E/I ratio and mean LFP metric across sessions and channels, and Spearman rank correlation was computed between mean E/I ratio and mean LFP metric. All statistics were evaluated at α=0.05.

