## Supplement for "Measuring excitation/inhibition balance through field potentials"

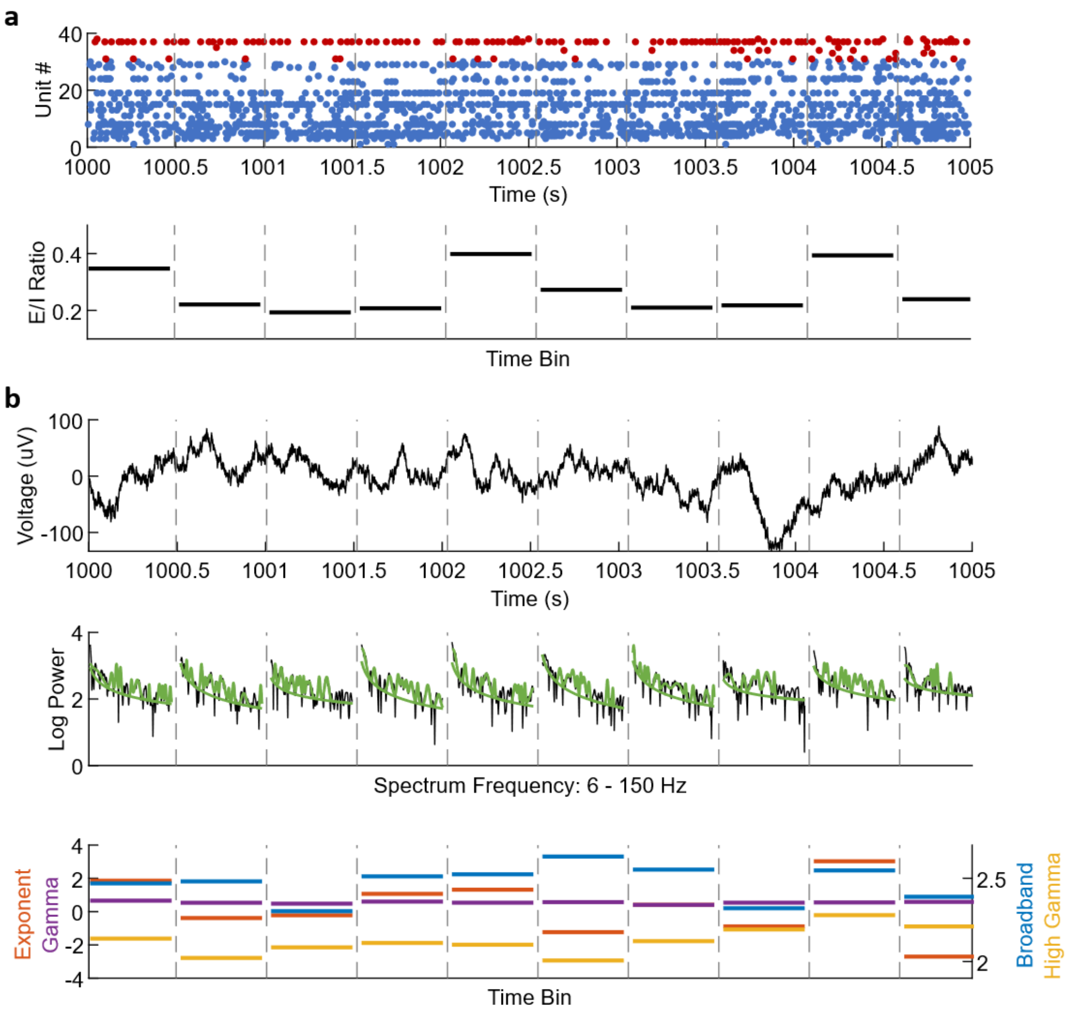


**Supplementary Figure 1. Analysis overview. (a)** Representative 5-second recording from medial prefrontal cortex showing spike rasters for principal cells (blue) and interneurons (red; top). The excitation/inhibition (E/I) ratio was calculated within consecutive 512 ms windows as the mean firing rate of principal cells divided by the mean firing rate of interneurons (bottom; dashed lines denote window boundaries). **(b)** Local field potential (LFP) recorded over the same interval (top), with power spectra computed for each 512 ms window (black) and fit with FOOOF (green; middle). Extracted spectral features (bottom) were subsequently correlated with the corresponding E/I ratios.

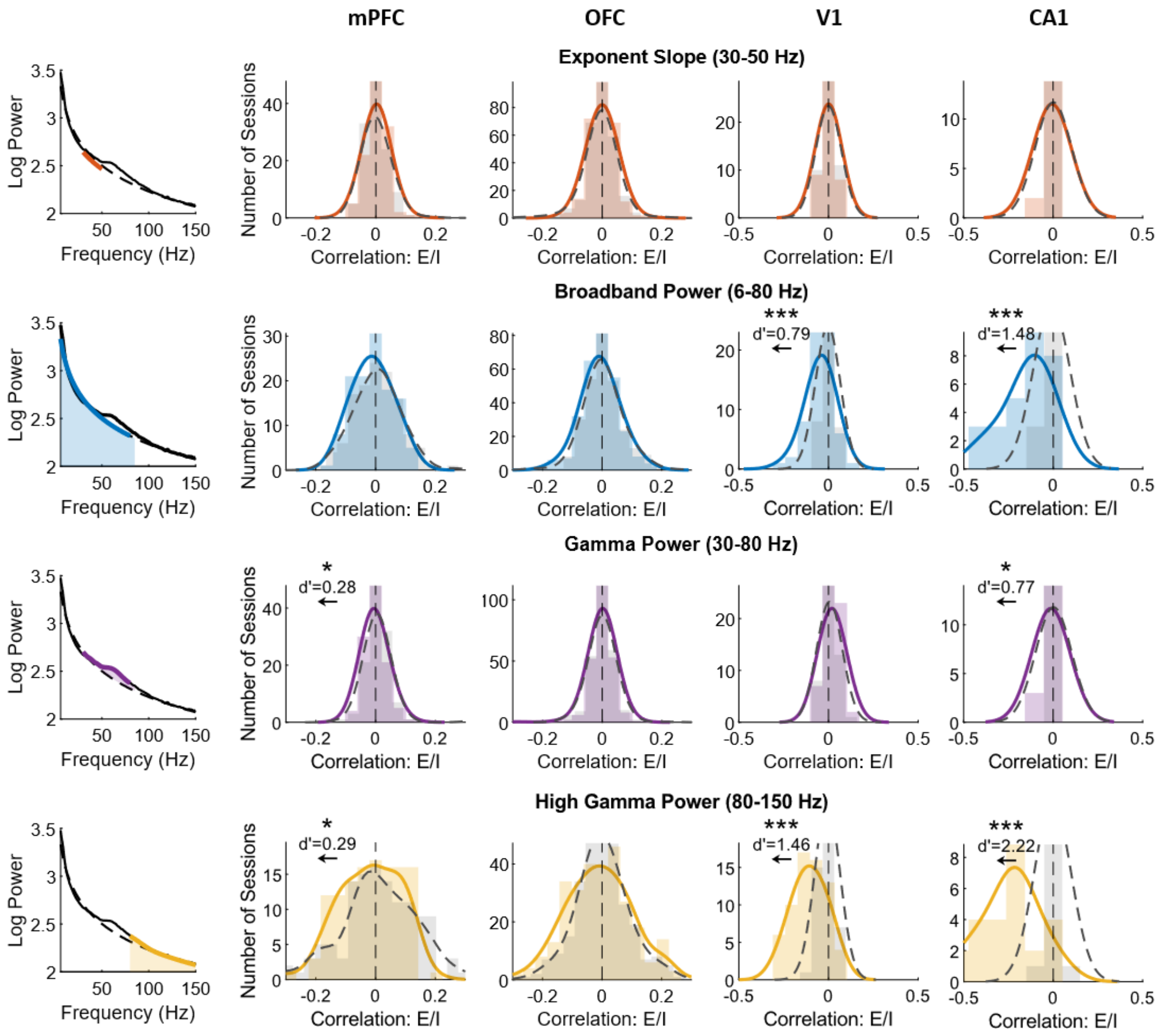


**Supplementary Figure 2. Local field potential metrics track excitation/inhibition balance across cell-type classification methods.** Relationship between local field potential (LFP) metrics (aperiodic exponent, broadband power, gamma power, high-gamma power) and the excitation/inhibition (E/I) ratio in medial prefrontal cortex (mPFC), primary visual cortex (V1) and hippocampal CA1 using cross-correlogram-based cell-type classification, and in orbitofrontal cortex (OFC) using spike width-based classification. Coloured histograms (solid curves) show real data; grey dashed curves show a shuffled null distribution. Note that cross-correlogram-based classification in CA1 may be confounded by synchronous population activity, which can obscure putative synaptic connections, and spike width-based classification in OFC was unreliable due to low spike waveform quality. *p<.05, **p<.01, ***p<.001.

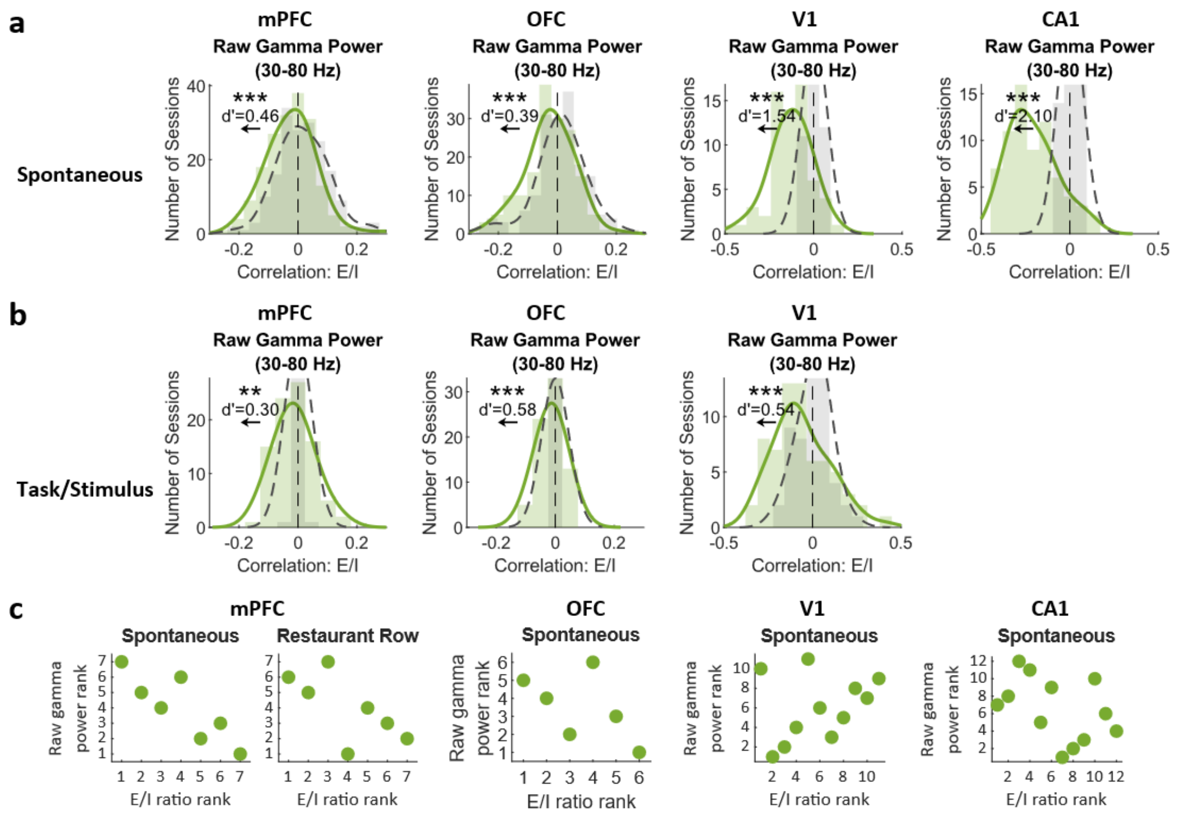


**Supplementary Figure 3. Raw gamma power correlates with excitation/inhibition balance.** (a) Distributions of correlation coefficients between raw gamma power (30-80 Hz) and the excitation/inhibition (E/I) ratio during spontaneous activity in medial prefrontal cortex (mPFC), orbitofrontal cortex (OFC), primary visual cortex (V1) and hippocampal CA1. **(b)** Distributions of correlation coefficients between raw gamma power (30-80 Hz) and E/I ratio in mPFC and OFC during the Restaurant Row decision-making task and V1 during drifting visual gratings. Coloured histograms (solid curves) show real data; grey dashed curves show a shuffled null distribution. *p<.05, **p<.01, ***p<.001. **(c)** Rank-order correspondence between animals' mean E/I ratio and raw gamma power across brain regions and task contexts. Each dot represents one animal. Animals are ranked from lowest to highest E/I ratio on the x-axis, and ranked from lowest to highest gamma power on the y-axis.

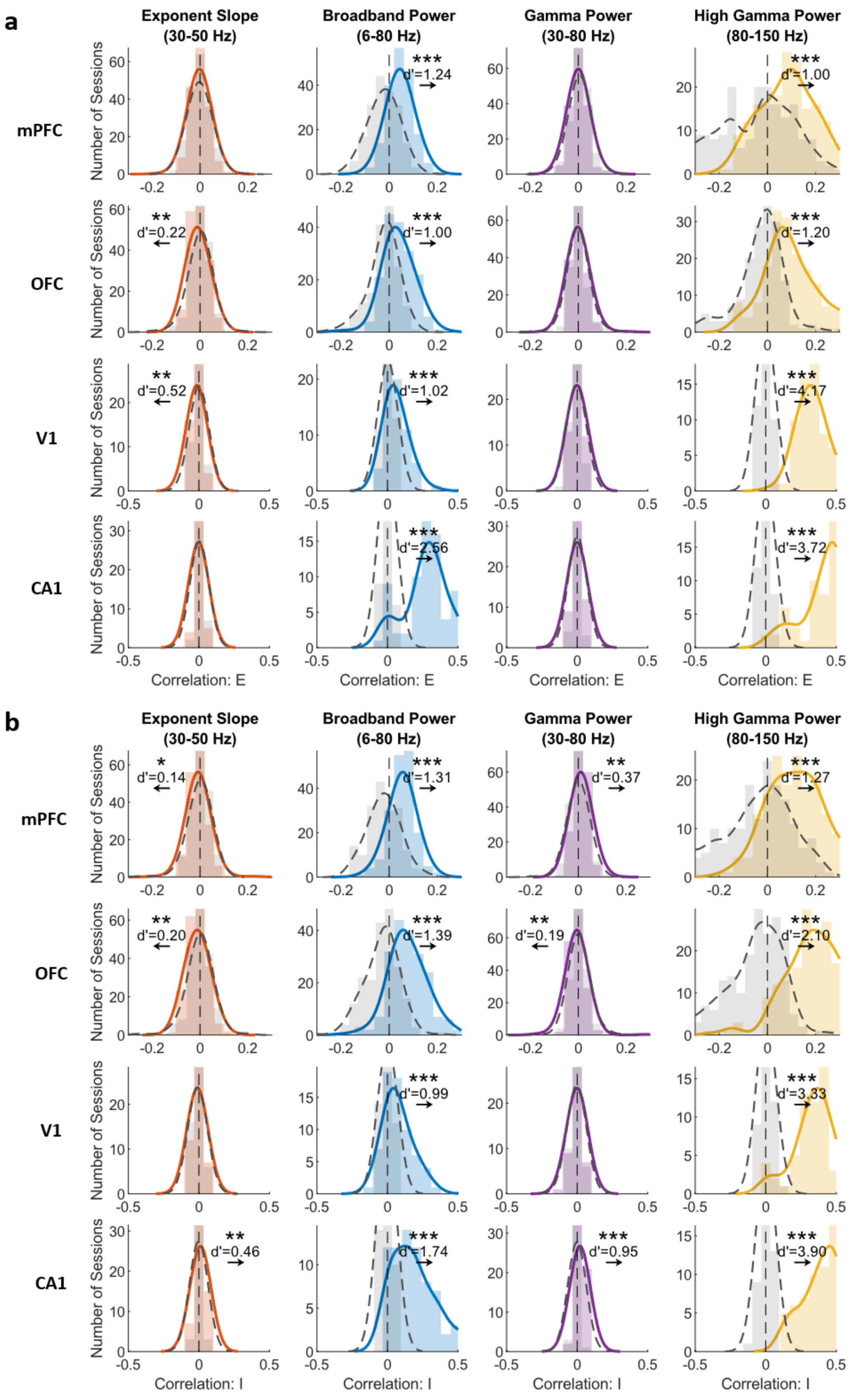


**Supplementary Figure 4. Local field potential metrics correlate with excitatory and inhibitory firing.** **(a)** Relationship between putative excitatory firing rates and each local field potential (LFP) metric (aperiodic exponent, broadband power, gamma power, high-gamma power) in medial prefrontal cortex (mPFC), orbitofrontal cortex (OFC), primary visual cortex (V1) and hippocampal CA1. Broadband power and high gamma power showed positive correlations with excitatory firing rates across mPFC, OFC, V1 and CA1 (Mann-Whitney U test; mPFC: Broadband z=10.25, p<.001, d'=1.24; High Gamma z=8.18, p<.001, d'=1.00; OFC: Broadband z=8.61, p<.001, d'=1.00; High Gamma z=10.40, p<.001, d'=1.20; V1: Broadband z=5.22, p<.001, d'=1.02; High Gamma z=9.89, p<.001, d'=4.17; CA1: Broadband z=8.72, p<.001, d'=2.56; High Gamma z=10.21, p<.001, d'=3.72). Coloured histograms (solid curves) show real data; grey dashed curves show a shuffled null distribution. **(b)** Same as in (a), but for putative inhibitory firing rates. Broadband power and high gamma power showed positive correlations with inhibitory firing rates across mPFC, OFC, V1 and CA1 (Mann-Whitney U test; mPFC: Broadband z=10.99, p<.001, d'=1.31; High Gamma z=10.34, p<.001, d'=1.27; OFC: Broadband z=12.50, p<.001, d'=1.39; High Gamma z=15.29, p<.001, d'=2.10; V1: Broadband z=5.16, p<.001, d'=0.99; High Gamma z=9.39, p<.001, d'=3.33; CA1: Broadband z=8.71, p<.001, d'=1.74; High Gamma z=10.35, p<.001, d'=3.90; Figure S4C). Note that the x-axis scale for mPFC and OFC differs from V1 and CA1. *p<.05, **p<.01, ***p<.001.

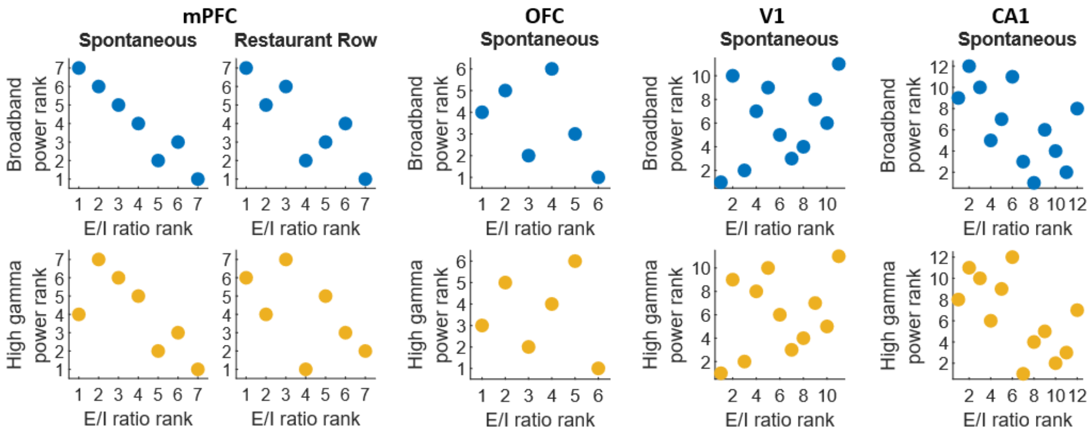


**Supplementary Figure 5. Rank-order correspondence between animals' mean excitation/inhibition ratio and local field potential metrics across brain regions and behavioural contexts.** Each dot represents one animal. Animals are ranked from lowest to highest E/I ratio on the x-axis, and ranked from lowest to highest broadband power (6-80 Hz; top row) or high-gamma power (80-150 Hz; bottom row) on the y-axis. mPFC, medial prefrontal cortex; OFC, orbitofrontal cortex; V1, primary visual cortex.

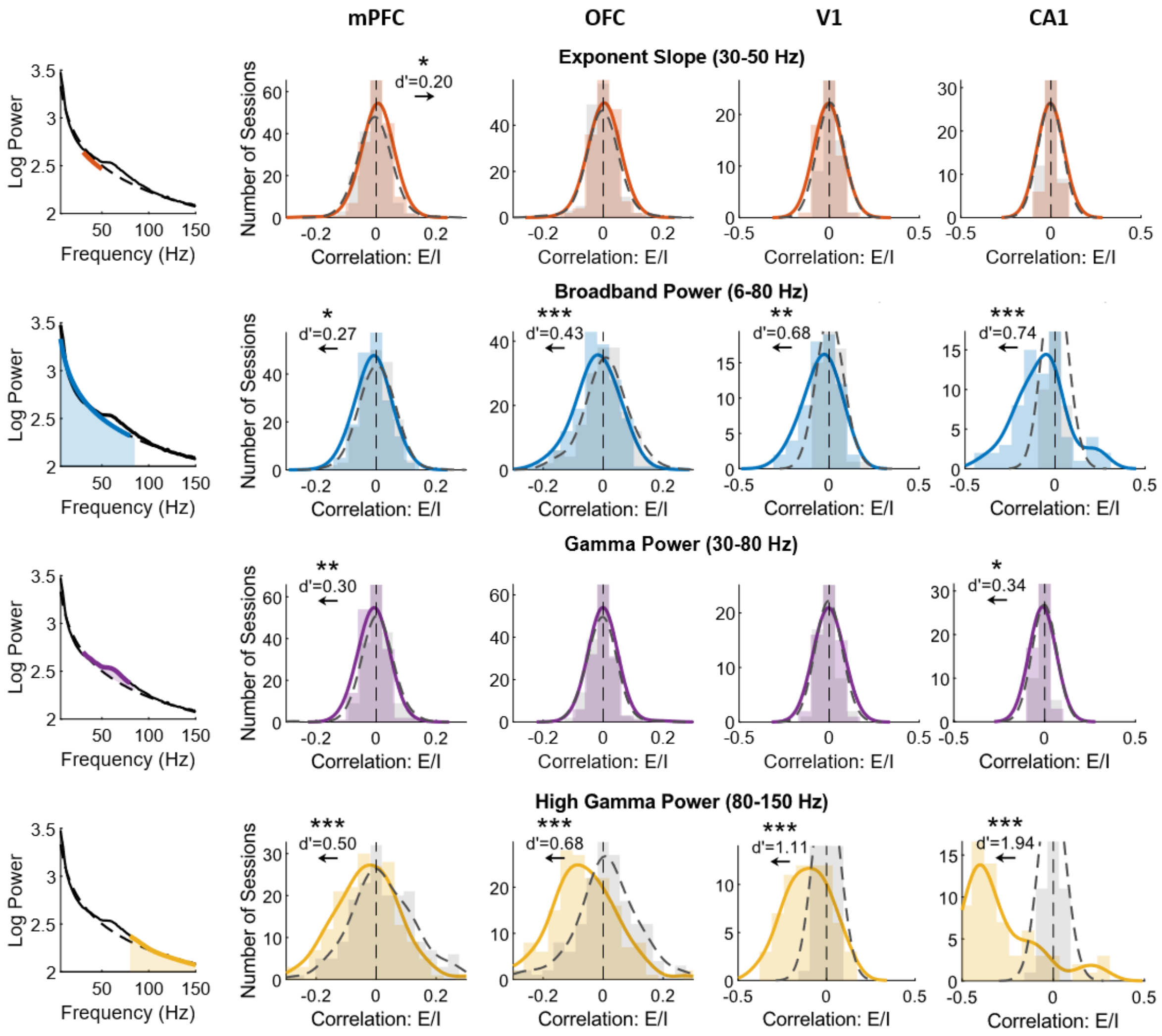


**Supplementary Figure 6. Correlations between local field potential (LFP) metrics and excitation/inhibition ratio in a single LFP channel.** Same as Figure 2, but correlation coefficients were computed using a single LFP channel rather than averaging across channels to confirm that results were not inflated by autocorrelation across channels. Coloured histograms (solid curves) show real data; grey dashed curves show a shuffled null distribution. Note that the x-axis scale for medial prefrontal cortex (mPFC) and orbitofrontal cortex (OFC) differs from primary visual cortex (V1) and hippocampal CA1. *p<.05, **p<.01, ***p<.001.
